# Evolutionary dynamics of specialization in herbivorous stick insects

**DOI:** 10.1101/367706

**Authors:** Larose Chloé, Rasmann Sergio, Schwander Tanja

**Affiliations:** Department of Ecology and Evolution, University of Lausanne, Switzerland; Institute of Biology, University of Neuchatel, Rue Emile-Argand 11, CH-2000 Neuchâtel, Switzerland

**Keywords:** Chaparral biome, host shift, plant-herbivore interaction, plant secondary metabolites, realized versus fundamental niche, redwood, *Timema* stick insect

## Abstract

Understanding the evolutionary dynamics underlying herbivorous insect mega-diversity requires investigating the ability of insects to shift and adapt to different host plants. Feeding experiments with nine related stick insect species revealed that insects retain the ability to use ancestral host plants after shifting to novel hosts, with host plant shifts generating fundamental feeding niche expansions. These expansions were not accompanied by expansions of the realized feeding niches however, as species on novel hosts are generally ecologically specialized. For shifts from angiosperm to chemically challenging conifer hosts, generalist fundamental feeding niches even evolved jointly with strong host plant specialization, indicating that host plant specialization is more likely driven by species interactions than by constraints imposed by plant chemistry. By coupling analyses of plant chemical compounds, fundamental and ecological feeding niches in multiple insect species, we provide novel insights into the evolutionary dynamics of host range expansion and contraction in herbivorous insects.

## Introduction

Long standing hypotheses suggest that the evolution of the tremendous diversity of insect herbivores (Gilbert 1979; Lawton 1983; Strong *et al*. 1984; Mitter *et al*. 1988; Farrell 1998; Novotny *et al*. 2006) relates to speciation driven by adaptation to novel host plants (Mitter *et al*. 1988; Schluter 2000; Dyer *et al*. 2007; Futuyma & Agrawal 2009; Matsubayashi *et al*. 2010; Hardy & Otto 2014). Many studies have focused on identifying the genetic basis of adaptations to novel hosts (Via 1991; Sezer & Butlin 1998; Feder *et al*. 2003; Nosil 2007; Soria-Carrasco *et al*. 2014; Simon *et al*. 2015), but what factors constrain the colonization of novel hosts at first remains largely unknown (Mayhew 2007; Winkler & Mitter 2008; Janz 2011). Indeed, multiple factors, including plant species-specific chemical compounds, which reduce insect growth and survival, are expected to hamper the ability of insect herbivores to shift to novel hosts (Scriber 1984; Hartley & Jones 1997; War *et al*. 2013a, b; Portman *et al*. 2015).

Overcoming constraints imposed by plant chemical compounds should be especially difficult for insect species that are specialized on few related host plant species, which appears to be the case for the vast majority of herbivorous insects (e.g., Fox & Morrow 1981; Scott 1986; Janzen 1988; Thompson 1994). Indeed, surveys of insect occurrences on plants in natural populations suggest that approximately 76% of all herbivorous insects are mono- or oligophagous, feeding on plant species belonging to a single genus or family (Forister *et al*. 2014). In spite of the widespread specialization, transitions from specialist to generalist habits have occurred repeatedly during the evolution of herbivorous insect clades (e.g., Funk & Bernays 2001; Nosil & Mooers 2005; Forister *et al*. 2012; Hardy & Otto 2014), questioning the idea that adaptation to plant chemical compounds generally hampers the colonization of novel hosts. Resolving this paradox has thus far been difficult because the majority of comparative and empirical studies on herbivore specialization (including the ones mentioned above) have only focused on the number of hosts used in natural population (i.e. the *realized* feeding niche; Colwell & Futuyma 1971; Futuyma & McCafferty 1990; Nyffeler & Sterling 1994; Blüthgen *et al*. 2006; Slatyer *et al*. 2013; Rasmann *et al*. 2014; Fordyce *et al*. 2016). Realized feeding niches depend on multiple factors, including insect adaptations to host plant chemistry, insect preferences (e.g., Dethier 1954; Forister *et al*. 2013) as well as species interactions (notably predation and competition; e.g., Hutchinson 1957; Novotny *et al*. 2006; Lewinsohn & Roslin 2008; Holt 2009; Ingram *et al*. 2012). However, little or no information is available on the range of plants allowing for survival, growth and reproduction of herbivorous insects (i.e., the *fundamental* feeding niche, Whittaker et al. 1973; Leibold 1995). Thus, the evolutionary dynamics of fundamental feeding niches are elusive and it even remains unknown whether the breadths of the fundamental and realized feeding niches generally change in parallel.

We hypothesized that the ability to use different plant species as hosts and consequently the breadth of the fundamental feeding niche is influenced by the evolutionary history of an insect lineage (see also Futuyma & McCafferty 1990). Specifically, if insect lineages can retain the ability to use their ancestral hosts as a food source after having shifted to a novel host, host shifts are expected to generate fundamental niche expansions (i.e., the lineage would become more generalist). By contrast, if insect lineages do not retain the ability to use their ancestral hosts, fundamental feeding niches will be independent of the evolutionary history of host plant use. More generally, colonization of novel host plants would be facilitated if insect lineages retained plasticity in host use present in their ancestors.

We used *Timema*, a small genus of herbivorous stick insects from western North America (Vickery 1993) to study the evolutionary dynamics of fundamental and realized feeding niches. Different *Timema* species have colonized plants from phylogenetically distant families, ranging from one to eight families of host plants per *Timema* species (Table 1). In terms of realized feeding niche, the *Timema* genus thus comprises a range of specialist to generalist species, and a tendency towards increased ecological specialization over evolutionary time was reported in a previous study (Crespi & Sandoval 2000). The genus originated about 30 million years ago (Riesch *et al*. 2017), in conjunction with the origin and spread of the chaparral biome to which most species are adapted (Sandoval *et al*. 1998; Crespi & Sandoval 2000). Ancestral *Timema* populations were most likely associated with angiosperms characterizing the chaparral biome, specifically the genera *Ceanothus* (lilac) and *Adenostoma* (chamise) (Sandoval *et al*. 1998; Crespi & Sandoval 2000). Nonetheless, transitions from angiosperm to conifer hosts have occurred multiple times in the genus. Ten of the 23 known *Timema* species regularly use conifers from one or multiple families as hosts (Table 1). At least two conifer species (redwood, *Sequoia sempervirens* and white fir, *Abies concolor*) represent recent host shifts, as both redwood and white fir are hosts for monophyletic groups of closely related *Timema* species (Fig. 1).

**Figure 1.**
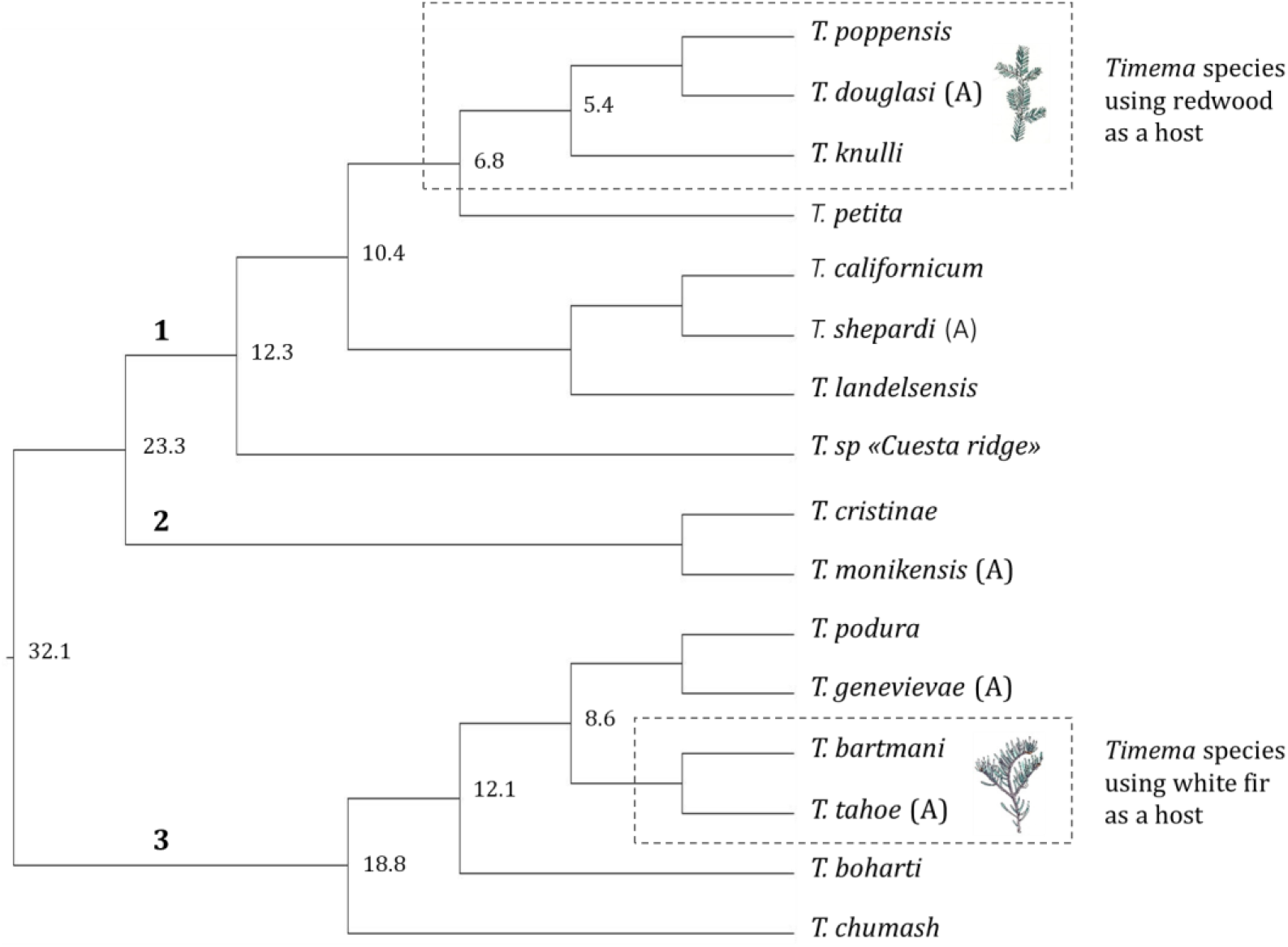
*Timema* phylogeny highlighting the species using the novel host plants redwood and white fir. Phylogeny redrawn from Riesch *et al*. 2017, with asexual lineages (A) added from Schwander *et al*. 2011. The phylogenetic position for the missing *Timema* species (see Table 1) is not known. Bold numbers 1, 2 and 3 correspond to the three described *Timema* clades, respectively Northern, Santa Barbara, and Southern clade. Node ages (Mya) are from Riesch *et al*. 2017.

**Table 1.**
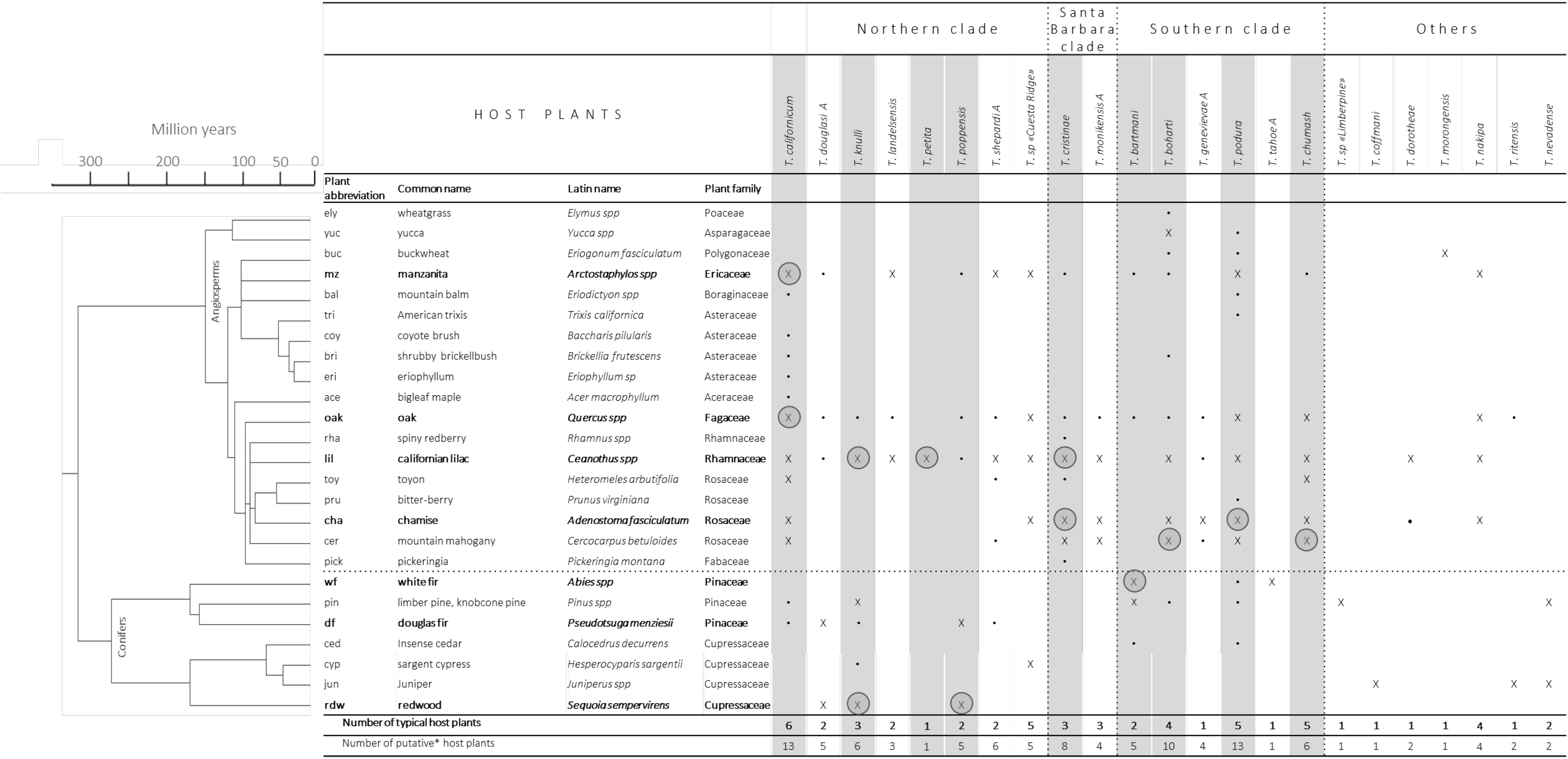
*Timema* species and their recorded host plants in the wild. Plants labeled with an “X” correspond to a common host for a given *Timema* species, where experimental evidence confirms that the plant is used as a food source. Plants labeled with “.” correspond to rare/anecdotal observations where it is unclear whether these plants are used as a food source (or solely for resting). Columns highlighted in gray indicate the *Timema* species used in the present study, sampling locations are specified in Table S1. Plants used for feeding experiments are written in bold. The plants on which the corresponding *Timema* populations were collected for this study are encircled. Note two of the *Timema* species are undescribed: *Timema* ‘Limberpine’, mentioned first by Sandoval & Crespi (2008), and *Timema* ‘Cuesta ridge’ from Riesch *et al*. (2017). The phylogenetic distances between the plant genera are estimated with information from the public database TIMETREE (http://timetree.org/; Hedges *et al*. 2015; Kumar *et al*. 2017).

Taking advantage of this variability in host plant use in *Timema*, we tested whether i) insect performance on host plants is constrained by plant phylogeny and plant chemical defenses, ii) the fundamental feeding niche breadth changes following a shift to a novel host, iii) insects retain the ability to use ancestral host plants following host shifts, and iv) fundamental and realized feeding niche sizes are correlated.

To characterize the realized feeding niches of the 23 known *Timema* species, we first generated a complete list of host plants for each species, using information from previous studies and field surveys. We then estimated the breadth of the fundamental feeding niche for nine of the 23 *Timema* species. To this end, we measured juvenile insect performance on seven phylogenetically diverse plants from the *Timema* host plant species pool (Table 1). This sampling strategy allowed us to study the evolutionary dynamics of specialization at the realized and fundamental niche levels. Finally, in order to explore potential mechanisms generating variable performances of insects on different plant species, we analyzed phenolic and terpenic secondary metabolites, which are toxins and/or feeding deterrents for many herbivorous insects (Bi & Felton 1995; Wink 1998; Acamovic & Brooker 2005; Dearing *et al*. 2005; Fürstenberg-Hägg *et al*. 2013).

## Material and Methods

### Realized feeding niches

In order to characterize the breadth of the realized feeding niche at the species level, we established a list of the host plants for each of the 23 known *Timema* stick insects species from the literature (Vickery 1993; Vickery & Sandoval 1997, 1999, 2001; Crespi & Sandoval 2000; Law & Crespi 2002; Sandoval & Crespi 2008; Riesch *et al*. 2017), and completed the list with personal observations (Table 1). We distinguished between plants for which we found evidence that *Timema* feed on them (hereafter “typical host plants”), and plants for which it was unclear whether they are used as a food source, or solely for resting (hereafter “putative host plants”; see Table. 1). In addition, we characterized the realized feeding niche at the population level for a subset of 22 populations from 9 species (between 1 and 6 populations per species; Table S1). To this end, we only chose locations where a minimum of 3 plants from the *Timema* host plants pool (Table. 1) were present. We then surveyed all these plants to determine the relative frequency of stick insects on each plant. (Table S1).

### Fundamental feeding niches

To measure insect performance on different hosts and their fundamental feeding niche breadths, we chose seven plants known to be commonly used by several *Timema* species, while trying to cover the phylogenetic diversity of all potential host plants (Fig. 1; Table 1). Stick insects for our experiments were collected from twelve populations belonging to nine *Timema* species throughout California (Table S1) using sweep nets. We only used fourth-instar juvenile females in order to minimize age-related effects, and to avoid the spurious effects of high mortality when manipulating younger instars. Between 10 and 80 females per host plant were used to measure survival and weight gain over 10 days, for a total of 70–220 females per population (1330 insects in total; see Fig. S1 for details on the experimental set-up). The large variation in numbers of insects per population was generated by the natural variation in the availability of forth instar females in different populations, as well as by the high mortality on certain plants that prevented us from obtaining weight gain estimates for all *Timema* populations. Whenever possible, we used more females for combinations generating high mortality.

### Evaluation of phylogenetic constraints regarding host use

We first tested whether closely related *Timema* species had similar performances (survival and weight gain) on the different plants. Branches from the most recent *Timema* phylogeny (Riesch *et al*. 2017) were pruned to create a phylogeny of the 12 populations from the nine species sampled for this study (Fig. 1). We used Mesquite 2.75 (Maddison & Maddison 2017) to reconstruct the ancestral states of the *Timema* performances on each of the seven plants (Mesquite module “Continuous-character Model Evaluation for phylogenetic signal testing”). Maximum parsimony with unordered, equal-weighted characters, and a cost of any state change = 1 was used to minimize the total number of character-state changes over the tree. We then compared the number of character-state changes inferred on the observed *Timema* phylogeny to the number of changes inferred on 1000 trees for which the characters were randomized across the tips in Mesquite. The null hypothesis that the character is randomly distributed on the phylogeny was rejected if the observed number of state changes fell outside of the upper or lower 5 percentiles of the random distribution (Maddison & Slatkin 1991).

### Estimations of the degree of specialization

To quantify the breadth of *Timema* feeding niches, we calculated the Tau specialization index (τ) (Yanai *et al*. 2004), as follows:

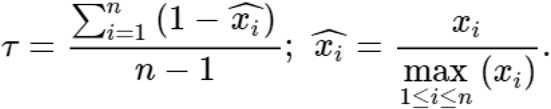

Where **n** corresponds to the number of plants, **x_i_** represents the frequency of occurrence (for the realized niche) or the weight gain (for the fundamental niche) on plant i, and **max (x_i_)** is the maximum occurrence or weight gain for the focal population. The index ranges from 0 (generalist) to 1 (pure specialist). We chose this measure to estimate the degree of specialization because of its robustness to small sample sizes and because our data were quantitative and continuous (Kryuchkova-Mostacci & Robinson-rechavi 2016). However, this index needs positive values to be calculated. We therefore transformed percentages of weight gain, which are negative when individuals lose weight, to relative weights of insects at the end of the feeding trials (i.e., an insect that lost 30% of its weight during the trial would be assigned the value 0.7, while one that gained 30% would be assigned 1.3). To test whether broad fundamental feeding niches translate into broad realized niches at the species or population level, we correlated the fundamental specialization indices Tau with the realized feeding niche breadths at the species and population levels, measured respectively by the number of host plants and the Tau indices based on the frequency of different host plants used within populations. We used Phylogenetic generalized least squares (PGLS) analyses to account for phylogenetic non-independence among *Timema* species. These analyses were conducted using the *ape* (Paradis *et al*. 2004) and *nlme* (Pinheiro *et al*. 2009) R packages (R Core Team 2017) using a Brownian motion model for trait evolution.

### Plant chemical profile characterization

We extracted and quantified compounds in the phenolic and terpene classes of secondary metabolites from leaves of the seven plant species included in our experiments (see Table 1), using methods adapted from Pratt *et al*. (2014) and Moreira *et al*. (2015). For each plant species, we extracted compounds from five independent replicates for both phenols and terpenes (see detailed methods for plant chemical analyses in Appendix S1).

To ordinate the chemical diversity data found across species, we conducted a principal component analysis (PCA) based on correlation matrices using the *FactoMineR* package in R (Husson *et al*. 2008). We tested whether plants have significantly different chemical compositions by estimating the chemical variation within and between species with a permutational multivariate analysis of variance (PERMANOVA) using 10.000 permutations with the *adonis* function (Anderson 2001) implemented in the R package *vegan* (Oksanen *et al*. 2007). We then tested for a correlation between the plant species phylogenetic distances and the chemical distances across the seven species tested using Mantel-tests with 10’000 permutations.

Finally, for the subset of chemical compounds that are present in multiple plants, we evaluated whether insect performances were negatively (or positively) correlated with the amount of a given compound. We conducted Spearman correlation tests (separately for each *Timema* population) between insect weight gain and each of the chemical compounds. These tests provided us for each *Timema* species with a list of chemical compounds significantly correlated to insect performance. We then tested whether these lists were more similar between different *Timema* populations than expected by chance, using hypergeometric tests with the *phyper* function in R (Johnson *et al*. 2005). Thus, we were not interested in the specific lists of significant chemical compounds per *Timema* population (which comprise many false positives due to multiple testing), but we were interested to see if the same compounds affect the performance of multiple *Timema* populations.

## Results

### Insect performances on different plants

The performance (survival and weight gain during 10 days) of *Timema* individuals was strongly dependent on the plant species tested. For ten of the twelve *Timema* populations, both survival and weight gain varied significantly among individuals reared on different plant species, while for the two remaining populations, only weight gain varied significantly (Table S2, Fig. S2). Insect survival and weight gain were also significantly correlated (Spearman rank correlation, r= 0.66, p < 0.0001), even though the most extreme situation (i.e., when all *Timema* of a given population died on a specific host plant before 10 days) could not be included in the analysis.

Generally, we found that insect performance was not maximal on the host plant they were collected on (henceforth referred to as the native host plant) (Table S2, Fig. S2). Indeed, for only five out of the 12 populations, individuals survived best on their native host plant, while for only six out of 12 populations they gained the most weight. In some cases, the performance of insects increased dramatically when individuals were reared on plant species they never use as host in the field. For example, 100% of *T. bartmani* survived for 10 days on lilac, while only 35.4% of them survived on their native host plant, white fir (Table S2).

We also observed that some host plant species are a consistently better food source than others. For instance, lilac was almost always the best food source, even for *Timema* species that never use lilac in natural conditions. Specifically, relative survival on lilac was high for all populations (between 76.9% and 100%, Table S2), and individuals from nine of the twelve *Timema* populations gained more weight when reared on lilac than when reared on any other plant species (Fig. S2). Lilac is the native host for only three of these nine populations (*T. cristinae*– lil, *T. knulli*-lil and *T. petita*), the six remaining ones were collected on manzanita (*T. californicum*-mz), chamise (*T. cristinae*-cha), oak (*T. californicum*–oak), mountain mahogany (*T. boharti* and *T. chumash*) or redwood (*T. knulli*-rdw). Only *T. podura, T. poppensis* and *T. bartmani* individuals had the highest weight gain when fed with their native host plant, with lilac ranking second.

Redwood was on the opposite end of the host plant quality spectrum, as it was only exploitable by *Timema* individuals originally collected on it. Relative survival on redwood for individuals from the two native redwood populations was high (75.0 and 86.7% for *T. poppensis* and *T. knulli*-rdw respectively; Table S2), while survival was low for all other *Timema* populations (ranging from 0% to 55.6%; Table S2). Similarly, *T. poppensis* and *T. knulli*-rdw were the only species that gained significant weight when fed with redwood for ten days (mean weight gain was 45.3% and 67.7% for the two species, respectively; Fig. S2). For the ten other populations, if individuals are able to survive for ten days on redwood, they typically lost weight (80% of surviving individuals) or only gained very little (20% of surviving individuals gained weight, with a maximum gain of 9.9%; Fig. S2). For the *T. bartmani, T. boharti, T. podura*, and *T. cristinae*-cha populations, not a single individual survived for ten days on redwood.

We observed the same pattern for *T. knulli*, the only *Timema* species using both redwood and lilac under natural conditions (Table 1). All individuals collected on redwood were able to live and grow on all tested plants (Table S2, Fig. S2). By contrast, practically all individuals of the same species collected on lilac died or lost significant weight on redwood (Table S2, Fig. S2).

### Degree of fundamental and realized specialization

The fundamental and realized feeding niche breadths were not correlated, neither at the species level, nor at the population level. At the species level, we found no significant correlation when considering the total number of host plant genera per *Timema* species (correlation corrected with Phylogenetic Generalized Least Squares (PGLS); r= −0.41, p= 0.43; Fig. 2), or when considering only the typical plant genera (PGLS; r= −0.17, p= 0.75)). The lack of correlation is unlikely caused by a lack of power as the general pattern is suggestive of a negative correlation between realized and fundamental niches rather than the expected positive correlation (Fig. 2). At the population level, we also found no correlation between Tau indices estimating the fundamental feeding niche and Tau indices estimating the realized niche (Pearson correlation test, r=0.02, p=0.91).

**Figure 2.**
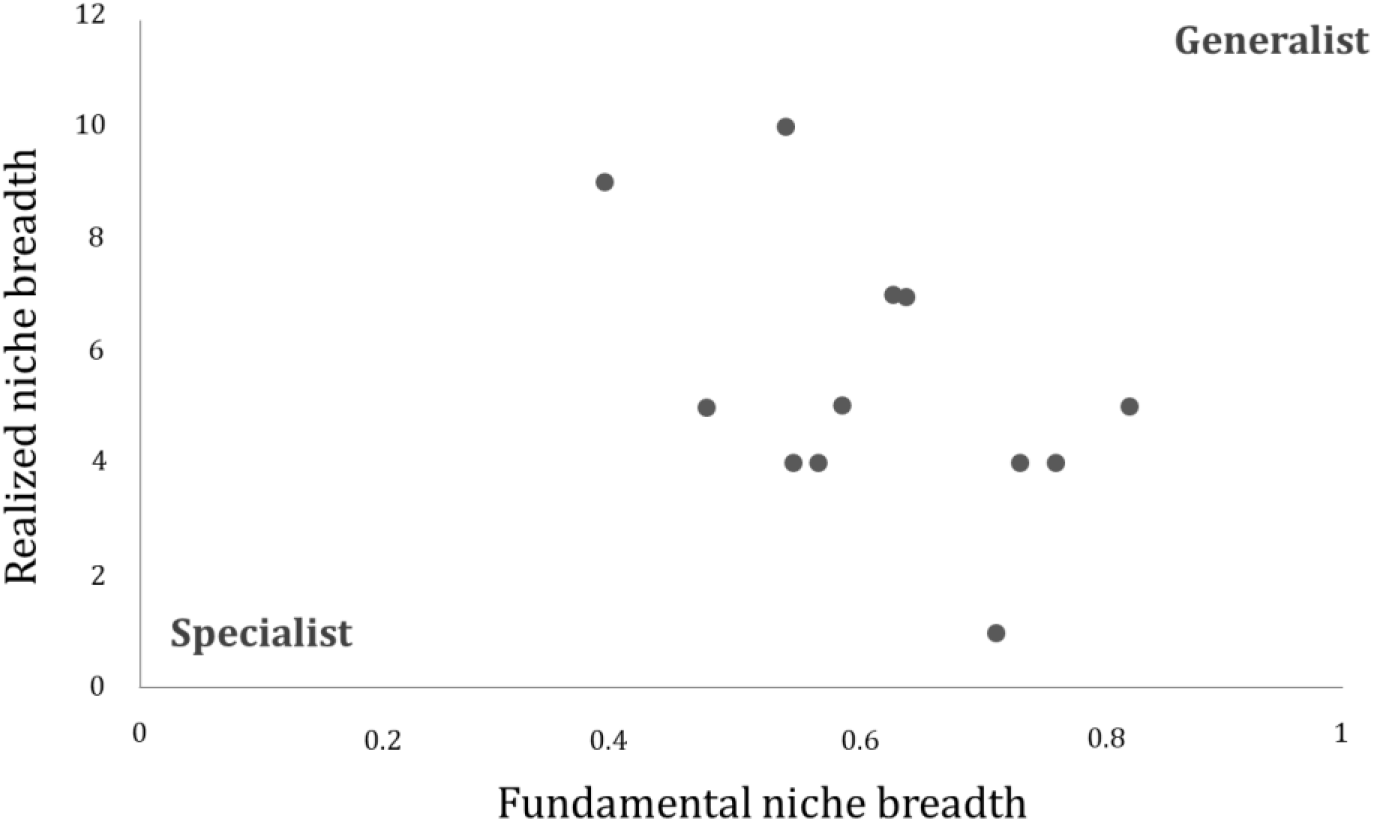
The size of the fundamental feeding niche does not constrain the realized feeding niche in *Timema*. Each point corresponds to a *Timema* population. For each population, the realized feeding niche breadth is estimated by the number of plant families used by the species, and the breadth of the fundamental feeding niche is estimated using the Tau index (based on insect weight gain).

The fundamental specialization indices showed that the two *Timema* species from redwood were the most generalist (Fig. 3A). The *T. knulli* population collected on redwood was also significantly more generalist (Tau = 0.23, 95% CI 0.19–0.30) than the population of the same species collected on lilac (Tau = 0.44, 95% CI 0.34–0.50). Hence, *Timema* native to redwood had a broader potential feeding niche than populations living on other host plants. In order to verify that this tendency was not only generated by the performance of the insects on redwood, we recalculated the Tau indices across six plants, excluding data from redwood. *T. poppensis. T. knulli*-rdw remained the most generalist species when the Tau indices were calculated without data from redwood (Fig. S5), and the Tau indices with and without redwood were strongly correlated (Pearson correlation; r: 0.96, p < 0.0001), indicating that the pattern was not solely driven by redwood.

**Figure 3.**
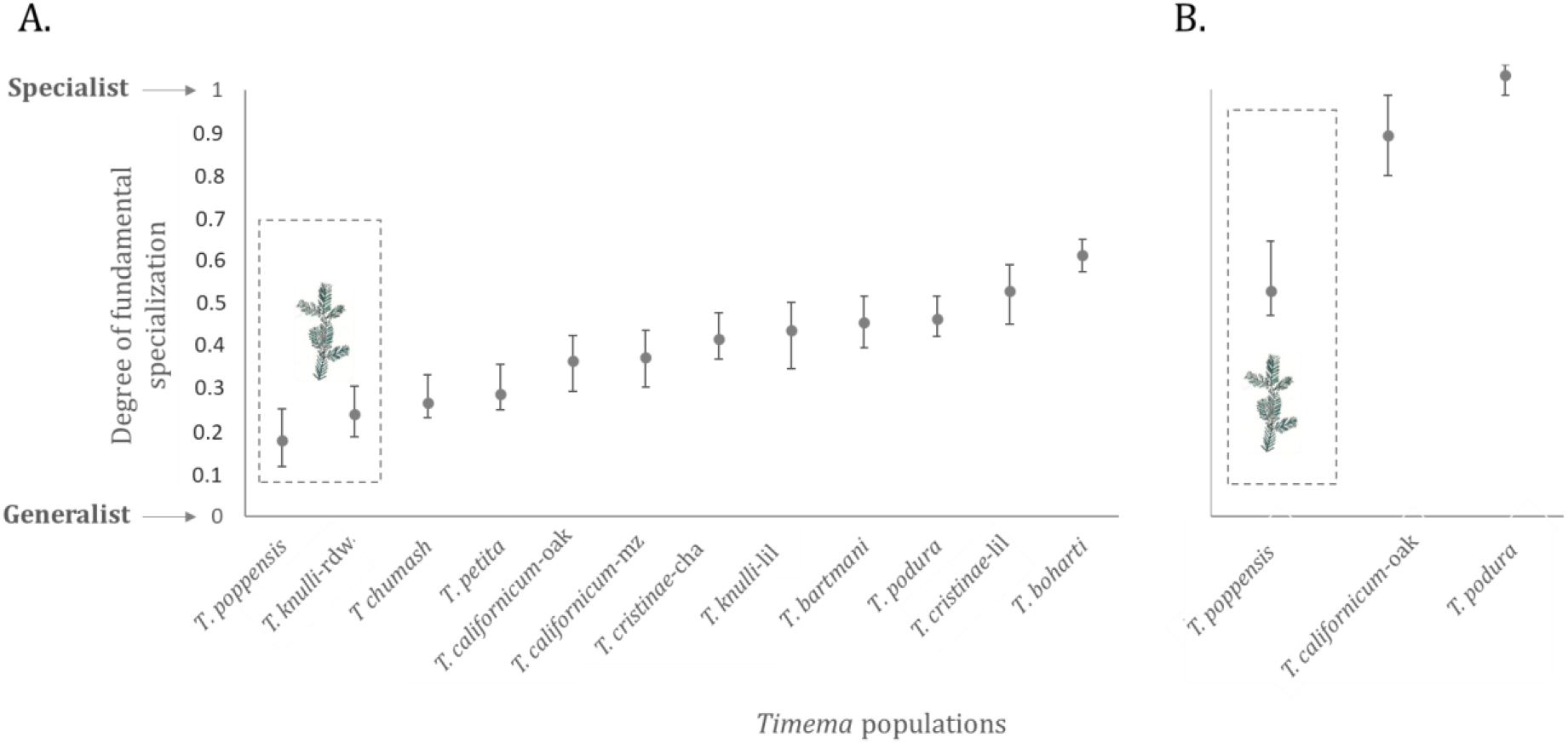
Breadth of the fundamental feeding niche of herbivorous stick insects. Niche breadth is quantified via the specificity index Tau (with 95% CI), based on insect weight gain on different plants (other measurements of specialization generate the same outcome, see Figures. S3, S4). The insect populations are listed from the least to the most specialist. Two independent analyses of specificity are presented. In the first one (A), the degree of specialization of twelve populations is based on their performance on seven plants from the *Timema* host plant pool. In the second one (B), the degree of specialization of a subset of populations is based on their performance on three novel plants not used by *Timema* stick insects in natural populations (sugar sumac, coyote bush and sage bush). The dotted rectangles highlight populations native to redwood.

These results suggest that the fundamental feeding niches of *T. poppensis* and *T. knulli*-rdw have expanded as a result of adaptation to redwood. To corroborate these findings, we reared individuals from three *Timema* species (*T. poppensis, T. californicum*-oak and *T. podura*) on plants not used as hosts by natural *Timema* populations (*Rhus ovata* (sugar sumac)*, Baccharis pilularis* (coyote bush) and *Artemisia californica* (sage bush)). Again, *T. poppensis* native to redwood performed better on these novel host plants than the two other insect species (Fig. 3B).

### *Effect of plant chemical composition on* Timema *performances*

To explore potential mechanisms generating variation in food quality among host plants, we studied the phenolic and terpenic secondary metabolites. We found a total of 521 different chemical compounds (28 phenols and 493 terpenes) across the seven plant species tested, with 84% of the variance explained by differences between species (PERMANOVA: *F*_6,28_ = 24.5, *p* < 0.001). In addition to chemical diversity, we also found that the total volume of compounds varied widely among plant species (volume measured as μg Gallic Acid Equivalent /g Dry Matter; average: 564μg/g; range 298 –1192), with a smaller volume in angiosperms (average: 310μg/g; range 298–331) than conifers (average: 902 μg/g; range 650–1192; Welch Two Sample t-test; t_2_ = –3.75; p= 0.063).

The PCA differentiated four plant groups, containing: 1) lilac, 2) oak, chamise, and manzanita, 3) redwood and douglas fir, and 4) white fir (Fig. S6). Distances between terpenic compositions of plants were correlated with the between plant phylogenetic distances (Mantel-test with 10.000 permutations, r = 0.77, p= 0.014), while there was no significant correlation for the phenolic compositions (Mantel-test with 10.000 permutations, r = –0.04, p= 0.47).

Most of the isolated terpenic and phenolic compounds were specific to a single plant or a subset of plants (Fig. S7). Specifically, 45.9% of the 521 compounds were detected only in a single plant, and only 1.5% of the compounds occurred in all seven plant species (Fig. S7). To test whether the performances of multiple *Timema* species were related to similar plant chemistries, we used the 162 compounds (31%) that occurred in at least three plant species. Among these, 84 (65 after FDR = 0.05 correction) were significantly correlated to insect weight gain in at least one *Timema* population. No single compound was found to be significantly correlated with the performance of *Timema* individuals collected from both angiosperms and conifers (Fig. 4). By contrast, 26 compounds (30.5%) were significantly correlated to the weight gain of insects from six of the nine populations living on angiosperms. One additional compound was further correlated to the weight gain of individuals of both populations collected from redwood (*T. poppensis* and *T. knulli-rdw*; Fig. 4). As phenols and terpenes are known to play an important role in plant defense against herbivorous insects, these compounds were expected to negatively affect insect performances. However, 59.2% of the compounds showed a positive effect (r varying between 0.77 and 0.99; Fig. 4), suggesting that some phenolic and terpenic compounds may favor rather than constrain *Timema* performance. The number of compounds significantly correlated to insect performance and shared among several populations significantly exceeded the amount of sharing expected by chance (Hypergeometric tests, p varying between 1e-06 and 1e-18).

**Figure 4.**
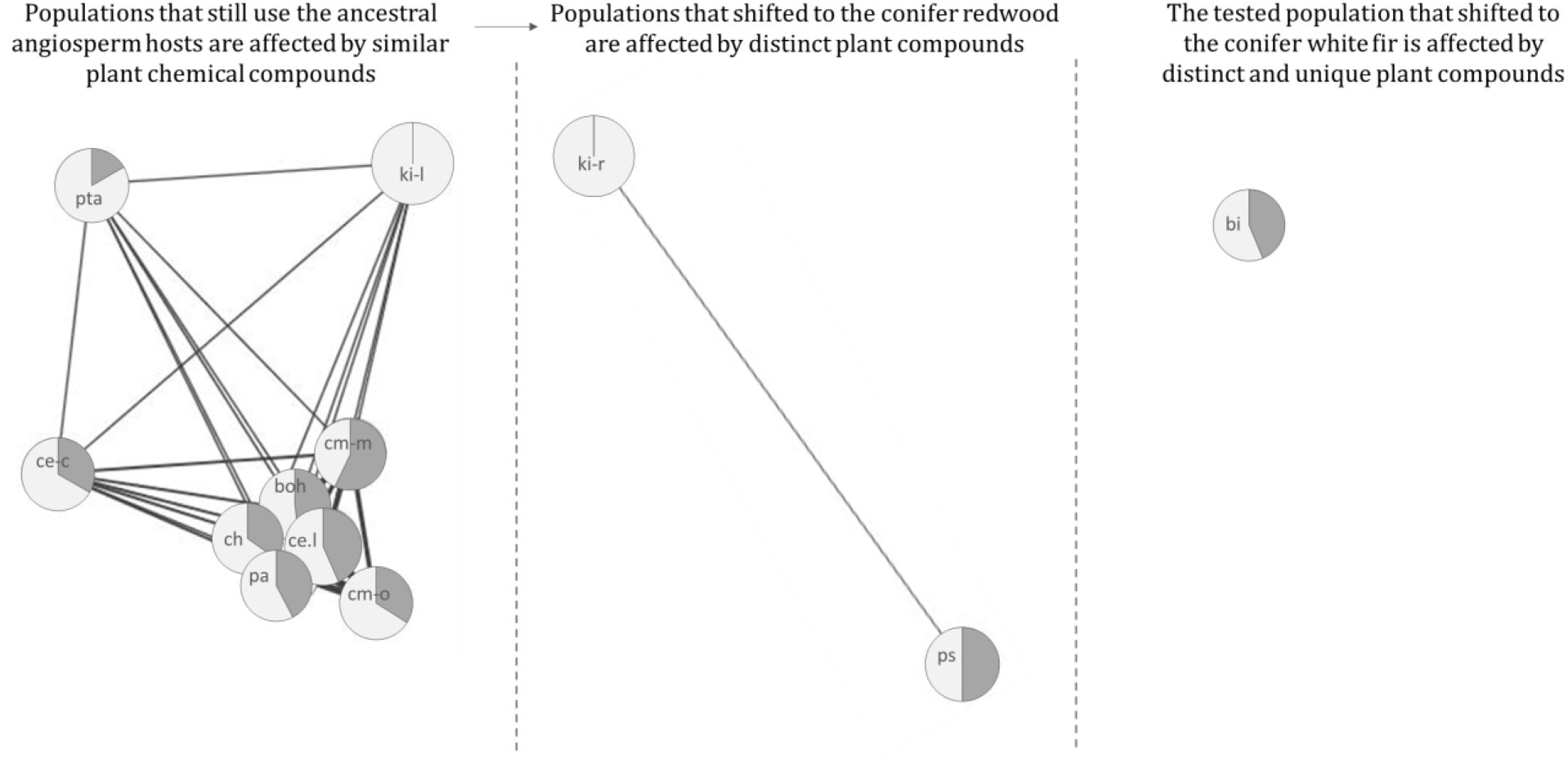
Similar plant chemical compounds affect the performance of insects native to angiosperm hosts, but different sets of compounds affect performances of insects native to conifers. Network built with Cytoscape 3.5.1 (Shannon *et al*. 2003). Circles in the network correspond to the twelve studied *Timema* populations. The length of the edges connecting two populations is negatively proportional to the number of shared compounds affecting insect weight gain (the more populations are affected by similar compounds the closer they are). The dashed lines separate groups of populations that are not affected by overlapping chemical compounds. *Timema* population name abbreviations are: bi: *T. bartmani* from white fir; boh: *T. boharti* from mahogany; cm-m: *T. californicum* from manzanita; cm-o: *T. californicum* from oak; ce-c: *T. cristinae* from chamise; ce-l: *T. cristinae* from lilac; ch: *T. chumash* from mahogany; ki-l: *T. knulli* from lilac; ki-r: *T. knulli* from redwood; pa: *T. podura* from chamise; ps: *T. poppensis* from redwood, pta: *T. petita* from lilac.

## Discussion

By studying the evolutionary dynamics of realized and fundamental feeding niches of multiple insect herbivores species in a phylogenetic framework, we developed novel insights into the mechanisms underlying feeding niche contractions and expansions. We analyzed the fundamental and realized feeding niches of *Timema* stick insects, which comprise a range of ecologically specialist to generalist species. We showed that insects expanded their fundamental feeding niches after shifting to new hosts. These fundamental niche size expansions occurred via two mechanisms. First, the species that shifted to novel hosts retained the ability to use plant groups used by their ancestors, even though the latest host shifts in *Timema* occurred 3–12 million years ago (Fig. 1). Second, adaptation to particularly toxic hosts (i.e., redwood) allows insects to metabolize chemically diverse plants, including plants currently not used as hosts by any species of the *Timema* genus. In combination, these mechanisms can explain how generalist insect herbivores (as measured from the realized feeding niche) can evolve from specialists, a pattern detected repeatedly at the macroevolutionary scale (Schluter 2000; Janz *et al*. 2001, 2006; Nosil & Mooers 2005; Stireman 2005; Winkler & Mitter 2008). Furthermore, fundamental feeding niche expansions following host shifts should facilitate future host shifts in the same lineage, which could generate frequent host turnovers via positive feedback loops of host adaptation and range expansion.

While several ecological factors, such as competition, predation or limited dispersal (e.g., Futuyma & Moreno 1988; Agosta 2006; Agosta & Klemens 2008) can drive ecological specialization, plant secondary chemistry has been brought forward as a key component driving insect performance and host plant specialization for herbivorous insects (e.g., Ehrlich & Raven 1964; Bi & Felton 1995; Dearing *et al*. 2005; Rosenthal & Berenbaum 2012; Portman *et al*. 2015). In the present study however, adaptation to a particular host plant chemistry does not explain ecological specialization in *Timema*. Indeed, the performance of *Timema* individuals was typically not maximized on their native host plant, as previously shown in feeding experiments with chamise and lilac for insect populations adapted to these two plants (e.g., Sandoval & Nosil 2005; Nosil 2007). We also found that *Timema* living on conifer hosts featured the broadest fundamental feeding niches of the genus, yet also the smallest realized one. In combination with the complete lack of correlation between fundamental and realized feeding niches in *Timema*, and the lack of phylogenetic constraint on fundamental niche size, these results suggest that plant secondary chemistry has little impact on insect host plant specialization. Accordingly, our analyses also revealed only minor effects of phenolic and terpenic compounds on insect performance.

Although we did not investigate the mechanisms driving host specialization in *Timema*, previous work in one species (*T. cristinae*) has shown that predation and plant preference (independently of plant quality) are key factors determining the distribution of insects on potential hosts (Sandoval 1994; Nosil *et al*. 2003; Sandoval & Nosil 2005). There is also accumulating evidence from herbivorous insects in general that preferences for host plant species are often not linked to the quality of plants as a food source, suggesting that insect preferences evolve more rapidly than insect physiologies (e.g., Rausher 1979; Thompson 1988; Valladares & Lawton 1991; Underwood 1994; Fritz *et al*. 2000; Faria & Fernandes 2001; Keeler & Chew 2008). Such preference-driven host plant selection in natural populations could help explain the lack of correlation between realized and fundamental niche size in *Timema*. Independently of the specific mechanisms driving host plant specialization in *Timema*, our results indicate that insect herbivores are more constrained by the biotic pressures of their environment than by their intrinsic physiological ability to metabolize particular plant species.

In the case of redwood, host plant chemistry might however indirectly mediate host plant use by relaxing insect-insect competition or pathogen pressure. Redwood is a host for only few herbivore species (Furniss 1977; Su & Tamashiro 1986; Grace & Yamamoto 1994), suggesting that competition on this host plant is low. In addition, laboratory experiments have shown that its wood inhibits the growth of bacteria (Scheffer 1966; Taha & Shakour 2016), and fungi (Shrimpton & Whitney 1968; Espinosa-Garcia & Langenheim 1990; Espinosa-Garcia *et al*. 1991), which may reduce pathogen pressure for insects. Finally, fires, being very common and an essential component of the Californian ecosystems (Minnich 1983; Brooks *et al*. 2004; Clinton *et al*. 2006), can favor redwood-insect associations. Thanks to their thick bark, redwoods can easily withstand high levels of burning (Jacobs *et al*. 1985; Ramage *et al*. 2010). *Timema* on redwood may thus survive fires while they would perish on more profitable hosts such as lilac or chamise. Using redwood may thus be overall beneficial even if it represents a non-optimal food source.

Our results suggest that the specific ability to use redwood is a key feeding innovation that allowed for range expansions in species that shifted to this host. Our feeding experiments showed that redwood is toxic to all *Timema* populations except for the native ones, while populations collected on redwood were able to survive and grow on all other tested host plants. Only three *Timema* species are known to use redwood in nature: *T. poppensis* and *T. knulli* (used in the present study), and *T. douglasi*, an asexual species very closely related to *T. poppensis* (Table 1). According to the most recent *Timema* phylogeny (Riesch *et al*. 2017), the last common ancestor of these three species occurred approximately 6.8 million years ago (Fig. 1), suggesting that the colonization of redwood happened around that time. The *Timema* genus appears to have originated in Southern California or Northern Mexico and expanded northward (Sandoval *et al*. 1998; Law & Crespi 2002), with several range expansion events for the species currently occurring at the northern end of the distribution such as *T. poppensis* and *T. douglasi* (the exact distribution of *T. knulli* is not known). Therefore, the incorporation of redwood in their diet was very likely of paramount importance for these herbivores to be able to expand their range northward. Indeed, the geographic distribution of redwood spreads over 750 km along the Pacific coast of the United States (Farjon 2005), while reaching further north than most other *Timema* host plants.

In conclusion, our study provides new insights into the consequences of host shifts for the breadth of the fundamental feeding niche. These consequences are highly relevant as they influence the probability for additional host shifts and potential host-associated diversification. Specifically, we showed that the ability to use ancestral hosts is maintained following major host shifts for at least 10 million years (as when moving from angiosperms to conifers), and that adaptations to chemically challenging hosts are not necessarily associated with decreased performance on alternative hosts. To the contrary, we here showed that adaptations to chemically challenging hosts allowed insects to metabolize a broad range of phylogenetically unrelated plants, including plants that have never been used as hosts in natural populations. More generally, the joint analysis of fundamental and realized feeding niches in multiple related insect species provides unique insights into the mechanisms driving the evolutionary dynamics of host range expansions and contractions in herbivorous insects.

## Acknowledgments

We thank Kirsten Jalvingh, Armand Yazdani and Bart Zijlstra for help in the field, Loren Bes (www.lorenbes.com) for the plant illustrations and Frédéric Bastian and Elsa Guillot for useful discussions regarding data analysis. We thank Jessica Purcell at UC Riverside for labspace and Darren Parker for proofreading. This study was supported by grant PP00P3 139013 of the Swiss FNS to TS and a fieldwork grant from the Swiss Zoological Society to CL.

## Supporting Information

## Appendix S1. Detailed methods for plant chemical profile characterization

We extracted and quantified compounds in the phenolic and terpene classes of secondary metabolites from leaves of the seven plant species included in our experiments (i.e., lil, cha, oak, mz, df, wf, rdw; see Table 1), using methods adapted from (Pratt *et al*. 2014) and from (Moreira *et al*. 2015), for terpenes and phenolics, respectively. For each plant species, we extracted compounds from five independent replicates for both phenols and terpenes. Leave samples for terpene extractions were stored in the freezer (−20°C) prior to use, while samples for phenol extractions were dried in an oven at 45°C for one week.

For phenol analyses, 100 mg of dried leaves per sample were reduced to powder with a pestle in liquid nitrogen, and phenols were extracted in 5 ml pure methanol (Sigma-Aldrich, CAS number 67-56-1). The methanolic solutions were kept at room temperature for 1 hour with continuous shaking. Thereafter, the extracts were sonicated for 10 minutes. Twenty-four hours later the tubes were centrifuged at 8000 rpm for 10 minutes and filtered. The collected supernatants were stored at 4°C until further use. Samples were analyzed by HPLC using a Grace C18 reversed phase column (3 μm, 150 × 4.6 mm; Grace Davison Discovery Science, Columbia, MD, USA) and an YL9100 instrument with diode array detection (YL Instrument Co., Anyang, Korea). The 15 μL injection was eluted at a constant flow of 0.7 mL min−1 with a gradient of acetonitrile and 0.25% phosphoric acid in water as follows: from 80% to 50% water in 5 min, then form 50% to 30% in 5 min, and kept at 30% for 7 min, and a final step from 30% to 5% in 4 min, followed by 5 min of equilibration time. Peaks were detected by a diode array detector at 270 nm (for hydrolizable tannins), 320 nm (for ferrulic acid derivates), 370 nm (for flavonoids) and 500 nm (for anthocyanins). Absorbance spectra were recorded from 200 to 900 nm. Peaks showing a characteristic absorption band of phenolics (Marbry *et al*. 1970) were recorded. Concentrations were calculated by using a standard curve that related peak areas to known gallic acid (for hydrolizable tannins), caffeic acid (for caffeic acid derivatives), quercitin (for flavonoids) and cyanidin (for anthocyanins) concentrations using 270 nm absorbance.

For terpene extractions, plant material was finely ground in liquid nitrogen and 250 mg were used for extraction in 2 mL n-hexane (Sigma-Aldrich, CAS number 110–54-3), with 20 □l internal standard (IS) added (tetraline; Sigma-Aldrich, CAS number: 119–64-2, 198 ng in 10 □l hexane). Five □l of each sample were subsequently injected into a GC-MS (Agilent 6890 Gas Chromatograph coupled with a 5973N Mass Selective Detector; Agilent, Santa Clara, CA, USA) fitted with a 30 m 9 0.25 mm 9 0.25 lm film thickness HP-5MS fused silica column (Agilent). We operated the GC in splitless mode with helium as the carrier gas (flow rate 1 ml min-1). The GC oven temperature program was: 1 min hold at 50°C, 10°C min-1 ramp to 130°C, 5°C min-1 ramp to 180°C, 20°C min-1 ramp to 230°C and 1 min hold at 300°C. We identified terpenes using Kovats retention index from published work (Loayza *et al*. 1995) and by comparison with commercial standards when available. We measured the richness (total number of compounds) and total production of individual compounds as a proportion to the IS.

---

**Table S1.** Sampled populations of nine *Timema* species. The number of individuals refers to the total number of individuals sampled in these locations on different host plants. For plant name abbreviations, see Table 1 in the main text.

| **Timema species** | **Location name (GPS coordinates)** | **Number of individuals per host plant sampled** |
| --- | --- | --- |
| *T. bartmani* | YMCA (34°09'48.8"N 116°54'22.6"W) | 0 oak, 0 pin, 350 wf |
| *T. boharti* | Sunrise (32°58'40.6"N 116°31'27.7"W) | 0 ad, 130 cer, 0 oak |
| *T. californicum* | Skyline (37°14'43.6"N 122°06'37.0"W) | 2 ad, 18 mz, 43 oak |
|  | Saratoga (37°11'47.0"N 122°02'27.1"W) | 4 ad, 12 mz, 4 oak |
|  | Summit (37°02'43.2"N 121°45'11.6"W) | 51 mz, 0 oak, 0 rdw |
| *T. chumash* | HW2_1 (34°15'42.4"N 118°06'27.6"W) | 45 cea, 70 cer, 250 oak |
|  | HW2_2 (34°16'12.5"N 118°10'06.8"W) | 18 ad, 5 cea, 11 oak |
| *T. cristinae* | Ojai1 (34°31'01.7"N 119°16'39.7"W) | 245 cea, 73 cer, 6 mz, 70 oak, 5 toy |
|  | Ojai2 (34°30'20.0"N 119°16'47.5"W) | 23cea, 62 cer, 11 mz, 28 oak |
|  | Ojai3 (34°31'59.6"N 119°14'51.8"W) | 8 ad, 2 cea, 20 cer, 8 oak |
|  | WTA1 (34°30'46.6"N 119°46'41.7"W) | 597 ad, 317 cer, 78 oak |
|  | WTA2 (34°30'22.3"N 119°46'05.3"W) | 81 ad, 1 cer, 8 mz, 9 oak, 2 toy |
|  | WTA3 (34°30'56.8"N 119°46'43.7"W) | 60 ad, 24 cea, 5 mz, 7 toy |
| *T. knulli* | HW1_1 (36°10'6.899"N 121°40'56.64"W) | 0 ad, 9 cea, 0 oak, 0 rdw |
|  | HW1_2 (36°14'50.8"N 121°46'54.4"W) | 0 cea, 0 oak, 13rdw |
|  | Big Creek (36°4'15.661"N 121°32'44.041"W) | 12 cea, 0 mz, 0 oak, 0 rdw |
| *T. petita* | HW1_3 (36°29'10.0"N 121°55'56.2"W) | 330 cea, 3 mz, 0 oak |
| *T. podura* | Indian (33°47'50.5"N 116°46'35.5"W) | 79 ad, 60 cea, 0 cer, 7 mz, 0 oak |
|  | Poppet (33°51'36.9"N 116°50'20.4"W) | 45 ad, 0 cea, 0 mz |
| *T. poppensis* | Fish_Rock (38°49'05.1"N 123°35'03.5"W) | 0 cea, 137 df, 14 rdw |
|  | Bear Creek (37°09'56.2"N 122°00'56.4"W) | 85 df, 0 oak, 35 rdw |
|  | Madonna (37°01'07.5"N 121°43'32.0"W) | 0 mz, 0 oak, 403 rdw |

**Table S2.** Relative survival of *Timema* individuals on different plants during ten days. For each *Timema* population, the survival on the native host plant is highlighted in grey. In the case of *T. boharti* and *T. chumash* the survival on their native host plant (*Cercocarpus betuloides*) is unknown as this plant was not included in the experiments. The proportion of deviance accounted for by the different plants in the GLMs was calculated using the modEva R package (Barbosa *et al*. 2013); Pearson’s chi-squared tests were performed to test whether plants explain a significant amount of deviance (p-value < 0.001: ***; < 0.01: **; < 0.05: *). For plant name abbreviations, see Table 1 in the main text.

| *Timema species* | Sample size per treatment | lil | cha | oak | mz | df | wf | rdw | % of deviance explained | p-value |
| --- | --- | --- | --- | --- | --- | --- | --- | --- | --- | --- |
| *T. bartmani* | 14 to 80 | 100.0 | 52.5 | 0.0 | 37.7 | 37,7 | 35.4 | 0.0 | 18.0 | 6.0e-07*** |
| *T. boharti* | 10 | 100.0 | 88.9 | 55.6 | 33.3 | 22.2 | 0.0 | 0.0 | 43.7 | 9.3e-06*** |
| *T. californicum-mz* | 10 | 90.0 | 100.0 | 100.0 | 90.0 | 70.0 | 50.0 | 10.0 | 42.1 | 2.4e-06*** |
| *T. californicum-oak* | 10 to 20 | 100.0 | 100.0 | 100.0 | 100.0 | 100.0 | 80.0 | 50.0 | 47.5 | 4.3e-04*** |
| *T. chumash* | 10 | 88.9 | 100.0 | 88.9 | 100.0 | 55.6 | 77.8 | 55.6 | 10.9 | 0.18 |
| *T. cristinae-cha* | 10 | 100.0 | 87.5 | 25.0 | 75.0 | 75.0 | 12.5 | 0.0 | 44.7 | 2.3e-05*** |
| *T. cristinae-lil* | 10 | 100.0 | 100.0 | 57.9 | 84.2 | 34.7 | 28.9 | 11.6 | 31.3 | 3.3e-04*** |
| *T. knulli-lil* | 10 to 20 | 100.0 | 100.0 | 26.7 | 93.3 | 26.7 | 26.7 | 6.7 | 29.7 | 3.9e-07*** |
| *T. knulli-rdw* | 24 | 90.5 | 86.7 | 71.4 | 77.5 | 100.0 | 82.1 | 86.7 | 3.9 | 0.36 |
| *T. petita* | 10 | 100.0 | 90.0 | 20.0 | 90.0 | 30.0 | 30.0 | 10.0 | 34.1 | 4.7e-05*** |
| *T. podura* | 15 | 76.9 | 100.0 | 23.1 | 92.3 | 23.1 | 7.7 | 0.0 | 41.1 | 4.7e-09*** |
| *T. poppensis* | 30 | 92.9 | 85.7 | 60.7 | 78.6 | 100 | 64.3 | 75.0 | 7.6 | 0.009** |

Barbosa, A.M., Brown, J.A. & Real, R. (2013). ModEvA–an R package for model evaluation and analysis.

**Figure S1.**
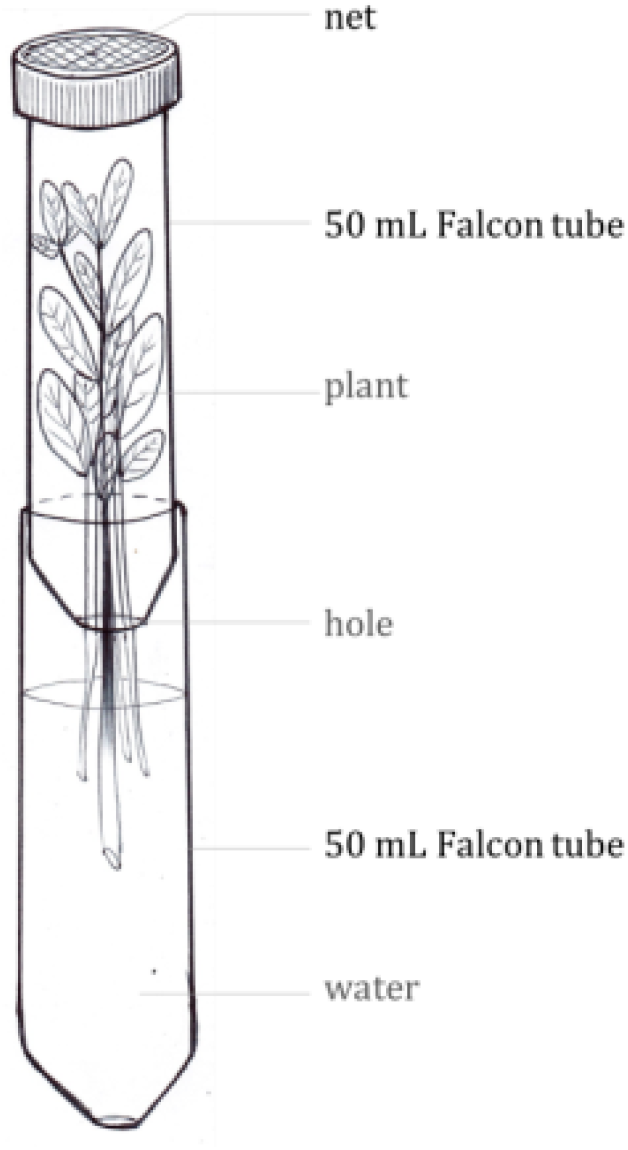
Illustration of the experimental system used to perform the feeding experiment. To measure the performance of insects on different plants, the collected juveniles were transferred to 50mL Falcon tubes containing a branch, with the broken end immersed in a water reservoir. Prior to the transfer, individual insects were weighed with an analytical balance (Kern ABT 120–5DM). During the ten days of the experiment, all tubes were observed daily to verify the survival of individuals and individuals that survived were weighted again at the end of the experiment.

**Figure S2.**
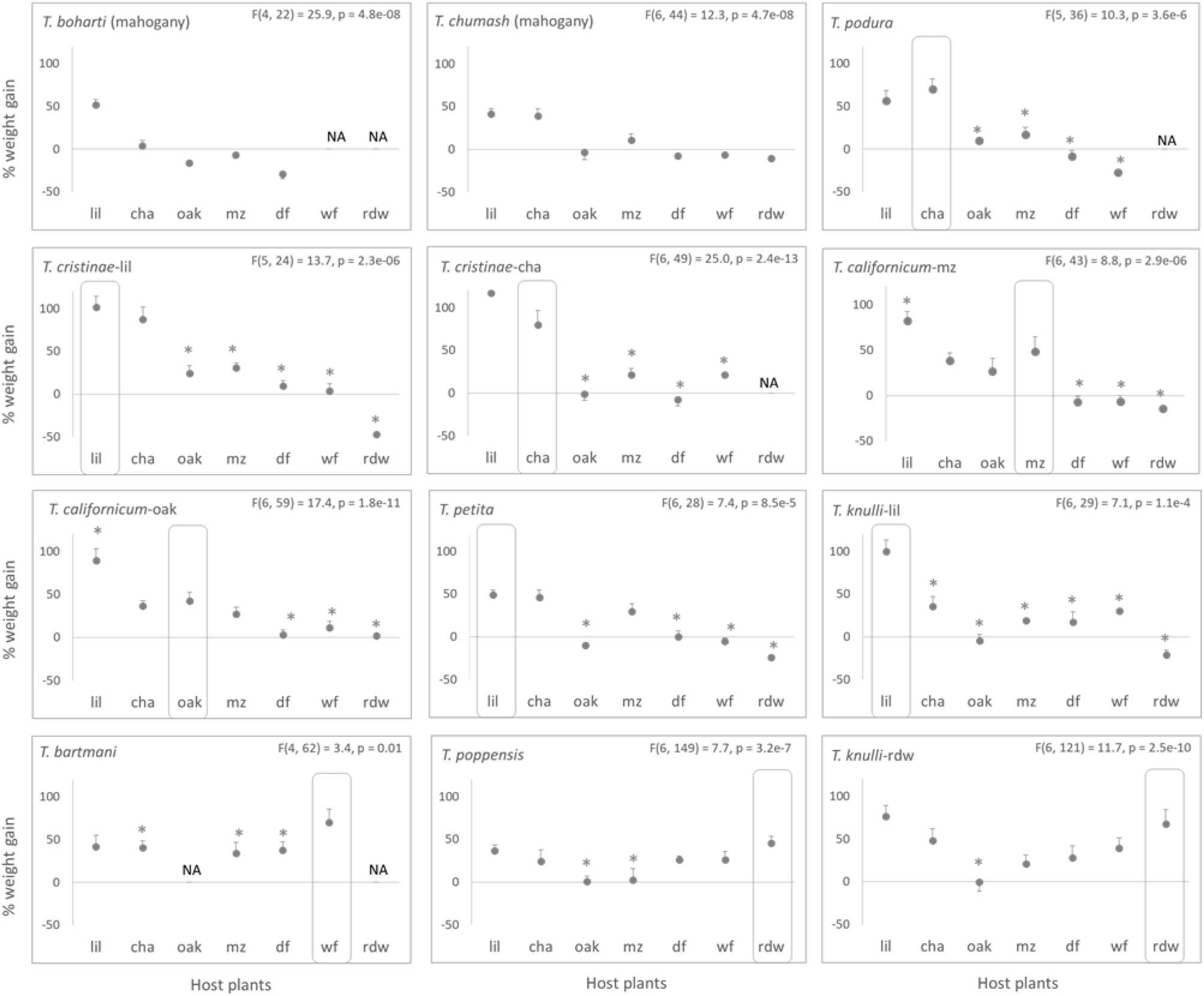
Percentages of weight gain for individuals fed with different plants for ten days. Each panel corresponds to a different *Timema* population, rectangles indicate native hosts. For each population, the amount of weight gained by individuals that survived during ten days on the different plants was compared using one-way ANOVAs. The asterisks indicate the plants on which the performance is significantly different from their performance on the native host (planned comparisons; * significant at p<0.05). For some plant by *Timema* population combinations, there are no weight gain data (NA) because all individuals died before the end of the experiment. For plant name abbreviations, see Table 1 in the main text.

**Figure S3.**
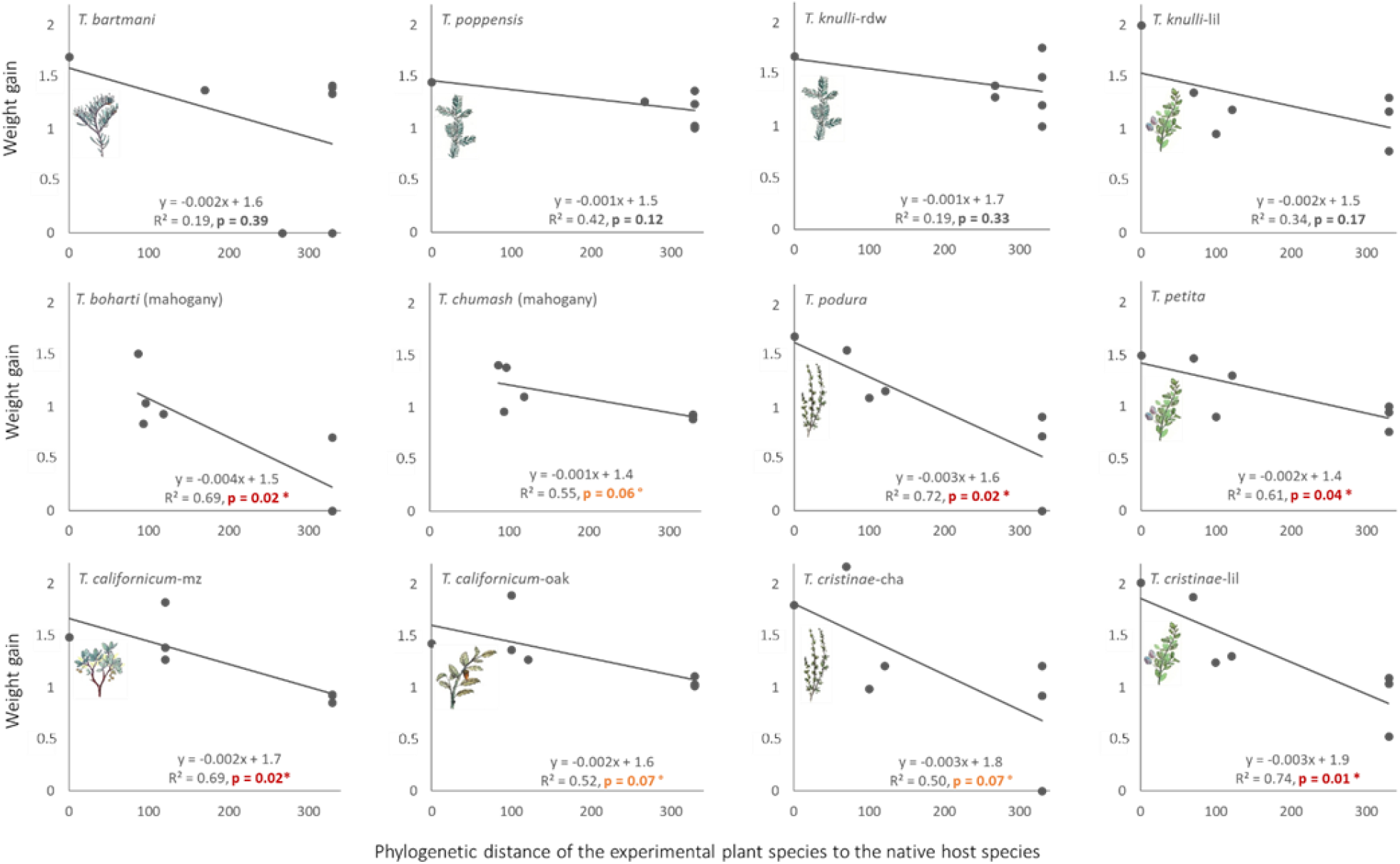
Relative weight gain of *Timema* individuals as a function of the phylogenetic distance between the native host plant and the plants used in the experiments. Native plants are indicated with icons (except for *T. boharti* and *T. chumash* where the performance on native hosts was not evaluated, see main text). Phylogenetic distances between plants are from Table 1 in the main text. Steeper slopes indicate more extensive specialization.

**Figure S4.**
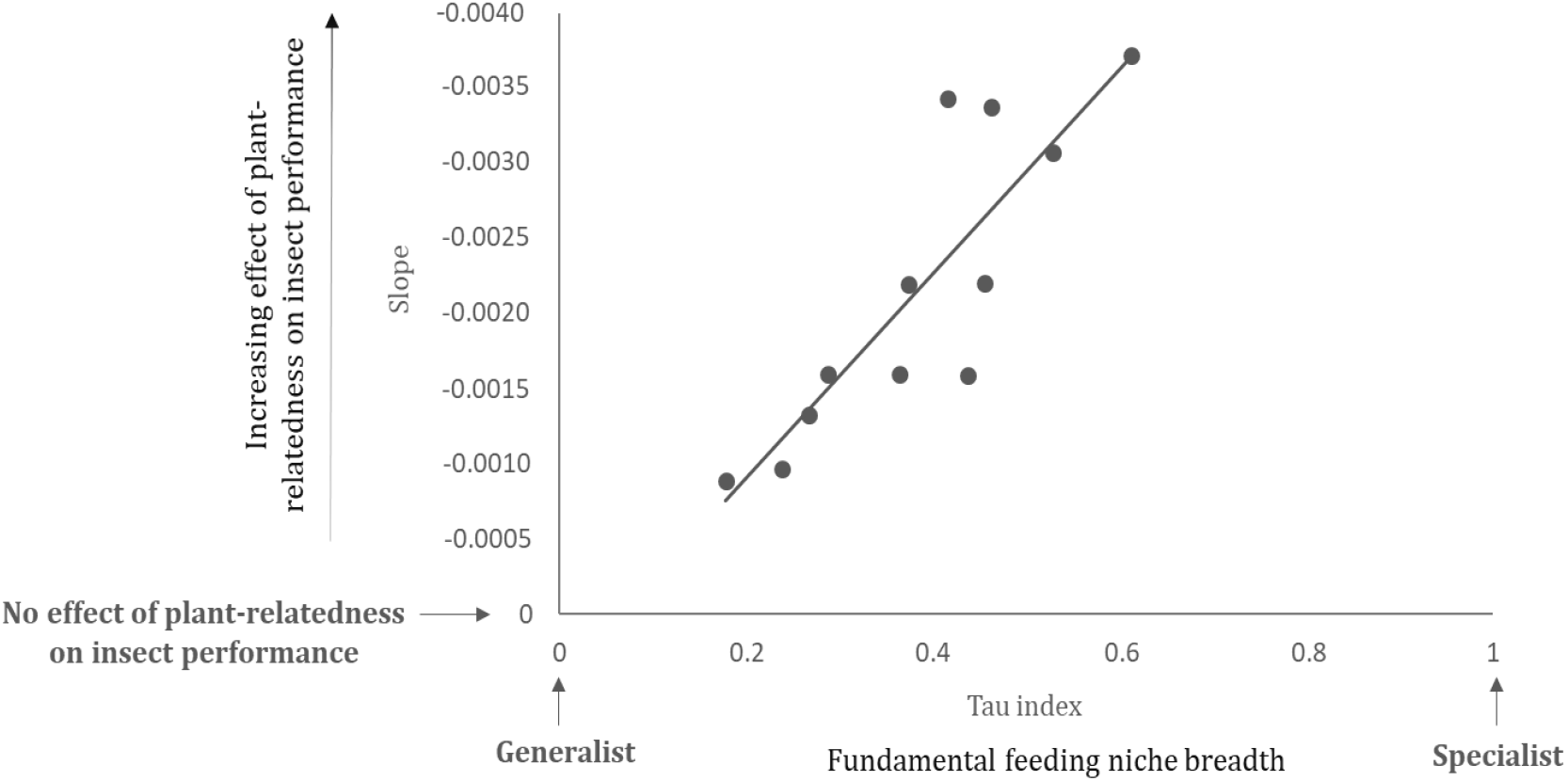
Correlation between two independent estimates of the degree of fundamental niche specialization. Each point corresponds to a *Timema* population. The Y-axis measures the performance decay of insects when fed with plants phylogenetically distant from the native host (slopes from Figure S5), the X-axis quantifies the specialization of insects via the Tau index. PGLS; r: –0.78, p= 0.025.

**Figure S5.**
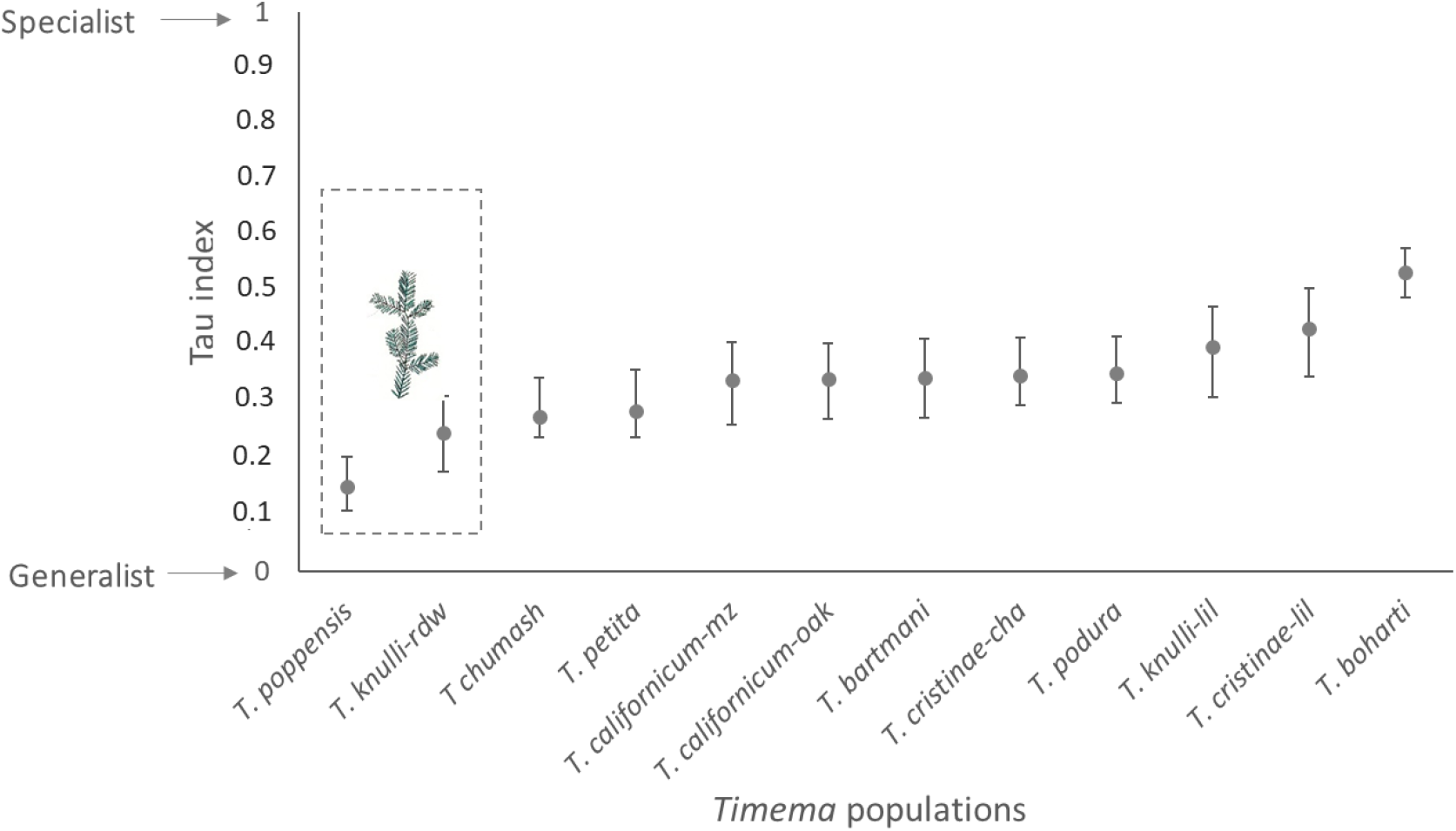
Breadth of the fundamental feeding niche of *Timema* from multiple populations. Niche breadth is quantified via the specificity index Tau (with 95% CI), based on insect weight gain on different plants (data from redwood excluded). The insect populations are listed from the least to the most specialist; *T. poppensis* and *T. knulli* native to redwood remain the most generalist populations even if data from redwood are excluded.

**Figure S6.**
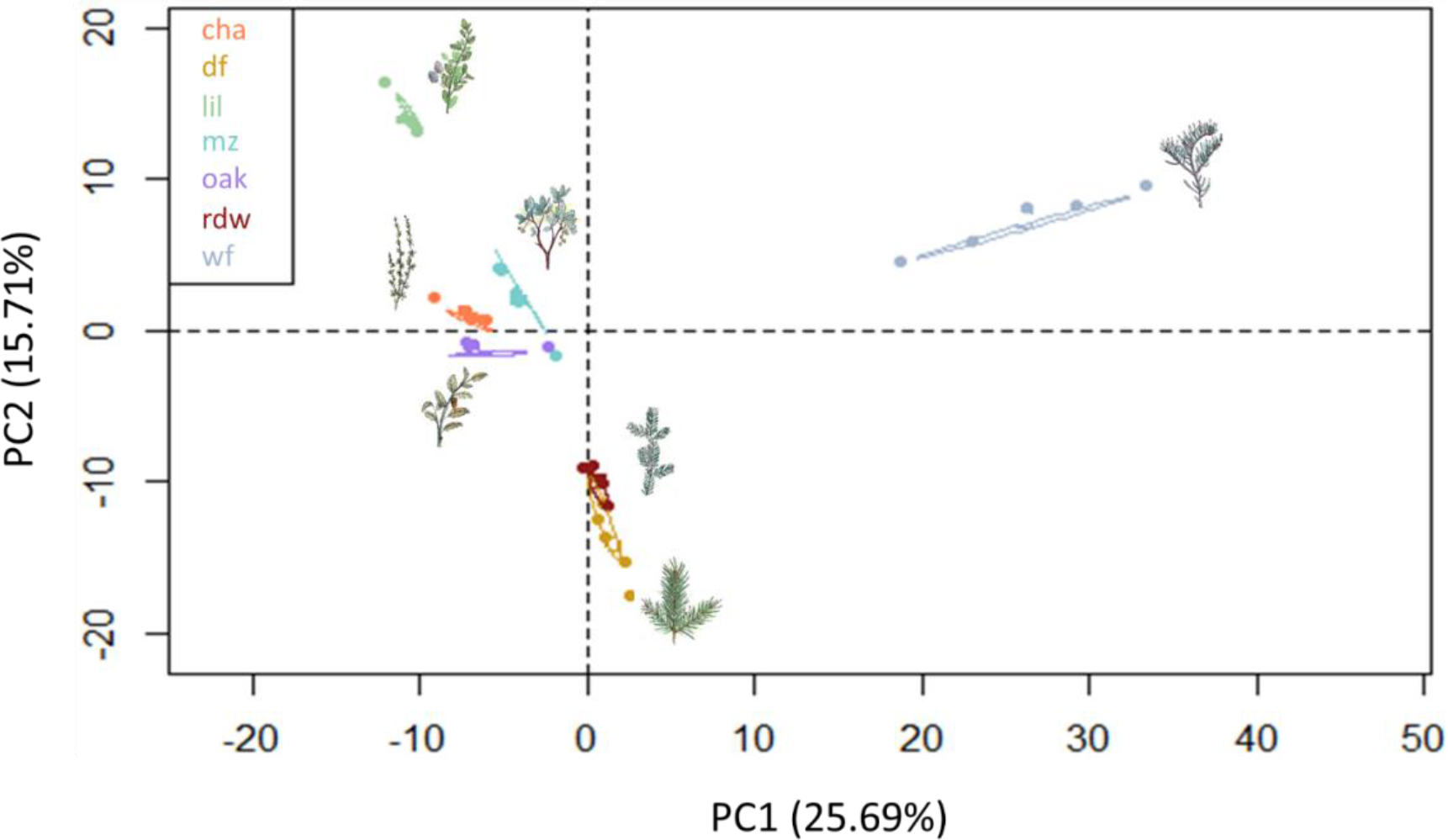
Principal component analysis based on the 521 plant chemical compounds (28 phenolic and 493 terpenic compounds). Percentages indicate the amount of variance explained by each axis. For plant name abbreviations, see Table 1 in the main text.

**Figure S7.**
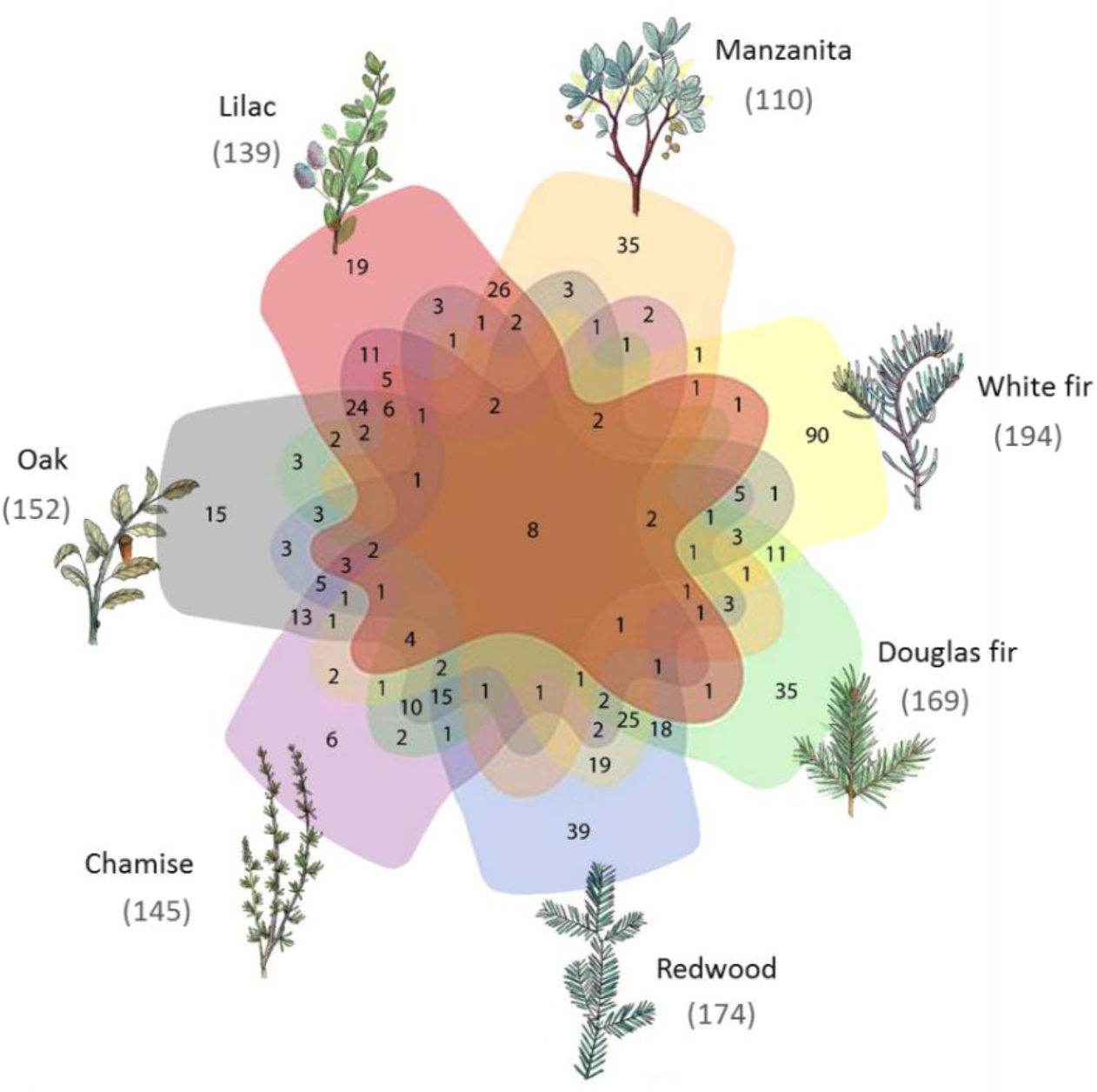
Specific and shared chemical compounds of different *Timema* host plants. The numbers in the Venn diagram indicate the number of terpenic and phenolic compounds shared among sets of plants and the number of species specific ones. Numbers in brackets indicate the total number of chemical compounds present in each plant.

